# Expression of human-specific SRGAP2C increases representational capacity in mice

**DOI:** 10.64898/2026.11.23.768640

**Authors:** Chiara S. Bramati, Jorge Ramirez-Ruiz, Franck Polleux, Graziella Di Cristo, R. Becket Ebitz, Roberto Araya

## Abstract

The brain of modern humans has evolved remarkable computational abilities enabling higher cognitive functions. Recent progress in evolutionary genomics and experimental neuroscience has identified several human-specific genetic modifiers of brain development that underlie the emergence of human-specific traits of cortical circuit connectivity and function. The human-specific gene *SRGAP2C* emerged by partial duplication of the ancestral gene *SRGAP2A,* which encodes a postsynaptic protein regulating synaptic development. SRGAP2C inhibits all known functions of SRGAP2A and its expression in mouse cortical pyramidal neurons leads to neotenic synaptic maturation, increased cortico-cortical connectivity, increased reliability of sensory coding by cortical layer 2/3 PNs, and improved learning of a sensory discrimination task. However, it remains unclear whether SRGAP2C expression in the mouse cortex is sufficient to produce changes in cognition. To test this idea, we trained adult wild-type (WT) and SRGAP2C mice on a Go-NoGo task that involves learning to associate a whisker stimulus with the presence of a water reward. SRGAP2C mice displayed a licking strategy that accelerated the learning of stimulus-reward associations at the cost of punishment timeouts compared to WT mice. The licking strategy was selective to the task, not driven by a general increase in consumptive behavior, and it accelerated the information that SRGAP2C mice gathered in early learning. SRGAP2C mice also displayed more structured behavior later in training and were, overall, better information channels than their WT counterparts. A computational model confirmed that SRGAP2C mice learning strategy increases representational capacity. Our results reveal that expression of human-specific gene SRGAP2C in cortical circuits is sufficient to increase the cognitive abilities of mice by optimizing the use of sensory information to guide action. The human-specific gene SRGAP2C might have contributed to the emergence of distinct functional cortical circuit features underlying our unique learning abilities and flexible decision-making.

## INTRODUCTION

Interacting with a complex and changing environment requires learning to associate complex and noisy sensory information with action choices and behavioral outcomes. Learning new associations depends on the reliable synthesis of sensory cues (*1*, *2*), internal states (*3*) and goals (*4–6*). In humans, the ability to learn associations is remarkably flexible and may play a role in our unusually rich behavioral repertoire (*7–10*). Indeed, the neocortex—a core structure underlying these computations—is elaborated in the primate lineage. The neocortex’s primary sensory and associative areas cooperate to enable the integration of sensory and contextual information with prior experience to produce learning (*11–15*). During human brain evolution, structural changes in the connectivity of cortical circuits allowed the emergence of human-specific cognitive capacities.

Human brain development is characterized by unique features among all mammals including great apes and other non-human primates such as a protracted developmental period (*16–18*), neotenic features of cortical development including synaptic development and maturation (*19–23*) and increased cortico-cortical connectivity (*17*, *18*). These human-specific features of brain development have been hypothesized to underlie the evolutionary emergence of unique cognitive traits such as increased cognitive abilities (*24*), more complex social interactions (*8*), the emergence of complex forms of language (*25*, *26*), and the influence of cultural transmission (*7*, *9*, *27*, *28*). However, establishing a causal relationship between human-specific features of cortical development and the emergence of our unique cognitive and social abilities has remained challenging. Even when we are able to compare different species in the same or similar cognitive tasks, it is essentially impossible to control for every possible difference in task, training procedure, experimental setup, not to mention controlling for prior exposure or adaptation to related tasks or training procedures (*29*). As a result, while modern humans may seem like exceptional problem solvers in some studies (*9*), humans can also appear to fall behind other species in other behavioral contexts and cognitive tasks (*30*, *31*).

Because of these challenges, we considered an alternative approach to comparative cognition research. Instead of comparing human cognition with other related species and speculating about underlying biological mechanisms, we genetically engineered the mouse genome to express a human-specific gene previously shown to implement human-specific developmental features of cortical connectivity, specifically in the cortex of mice throughout development in order to determine how it changes their cognitive capacities of mice compared to wild-type control mice. Specifically, our focus is on a human-specific gene linked to the maturation of cortical connectivity: SLIT-ROBO Rho GTPase Activating Protein 2C (SRGAP2C). SRGAP2C is a human-specific paralog of the mammalian gene SRGAP2, which is highly expressed during brain development (*32*, *33*). SRGAP2 promotes excitatory (E) and inhibitory (I) synaptic maturation and limits the number of both E and I synapses received by mouse cortical pyramidal neurons (*34*, *35*). The ancestral SRGAP2 (renamed SRGAP2A in humans) was duplicated at different timepoints of human brain evolution, which led to the appearance of the human-specific paralogs SRGAP2B and SRGAP2C approximately ¾ and 2.4 million years ago (mya) i.e. at the birth of the *Homo* lineage (*36*, *37*). In particular SRGAP2C, has been shown to partially inhibit all known functions of SRGAP2A and phenocopies a partial loss of function of SRGAP2A (*34*) by inducing SRGAP2A proteasome-dependent degradation to ∼50% of wild-type levels (*34*, *35*, *38*). Transgenic expression of SRGAP2C in mouse cortical pyramidal neurons is sufficient to delay synaptic maturation, inducing neotenic features in cortical pyramidal neurons during development and increased cortico-cortical connectivity in adult layer 2/3 pyramidal neurons (L2/3 PNs) (*34*, *35*, *39*) —producing some of the same features of human neocortical pyramidal neurons (*19–21*, *23*). Most importantly, it was recently shown that downregulation of human-specific *SRGAP2C* in human pyramidal neurons xenotransplanted in the cortex of mouse neonates leads to striking acceleration of synaptic development and functional synaptic maturation of these human neurons *in vivo* (*40*). These results demonstrate that the human-specific gene SRGAP2C is necessary and sufficient to induce human-specific features of synaptic development and cortical circuit connectivity.

Prior work investigating the behavioral impact of SRGAP2C-expressing mice suggested improvement in the ability of these mice to learned a challenging sensory discrimination task (*39*, *41*). However, these differences could be due to a variety of cognitive mechanisms, including changes at the level of perception, sensorimotor association, or even in the higher-level cognitive strategies used to learn perceptual associative tasks (*42*, *43*). To determine if and how SRGAP2C expression in cortical pyramidal neurons alters cognition, we trained wild-type (WT) and Nex^Cre+^SRGAP2C^F/+^ (SRGAP2C) mice in an active Go/NoGo detection task and compared their behavioral performance and cognitive strategies throughout task acquisition. Despite showing similar perceptual sensitivity levels (d’), we found that the two genotypes differed significantly in the strategies they used to solve the task. While both genotypes were able to learn the task, SRGAP2C mice accumulated more task-relevant information early in training and exhibited more structured behavior in later stages. Overall, they appeared to be more information-preserving than WT mice. We also leveraged a computational approach, based on a reinforcement learning model of our detection task, and found that increasing the representational capacity of the model is sufficient to reproduce all major features of SRGAP2C mouse behavior. These results suggest that the ability of SRGAP2C to modify cortical circuit connectivity and cortical circuit function (*39*, *41*) links the evolutionary emergence of this human-specific gene to the emergence of a human capacity for rich representational structures and to increased cognitive abilities by optimizing the use sensory information to guide action.

## RESULTS

### WT and SRGAP2C mice reach similar sensitivity levels in an active Go/NoGo sensory detection task

The Nex^Cre+^SRGAP2C^F/+^ transgenic mouse line (SRGAP2C mice) expresses the human-specific gene SRGAP2C in all excitatory pyramidal neurons of the neocortex and hippocampus in a Cre-dependent manner (*39*) (**Fig 1A**). We validated that the transgene leads to expression of SRGAP2C-HA only in Nex^Cre+^SRGAP2C^F/+^mice but not in SRGAP2C ^F/+^ (control) mice by performing anti-HA immunofluorescent staining on cryostat sections of the barrel cortex of adult Nex^Cre-^SRGAP2C^F/+^ and Nex^Cre+^SRGAP2C ^F/+^ mice (**Fig 1B**).

**Figure 1.**
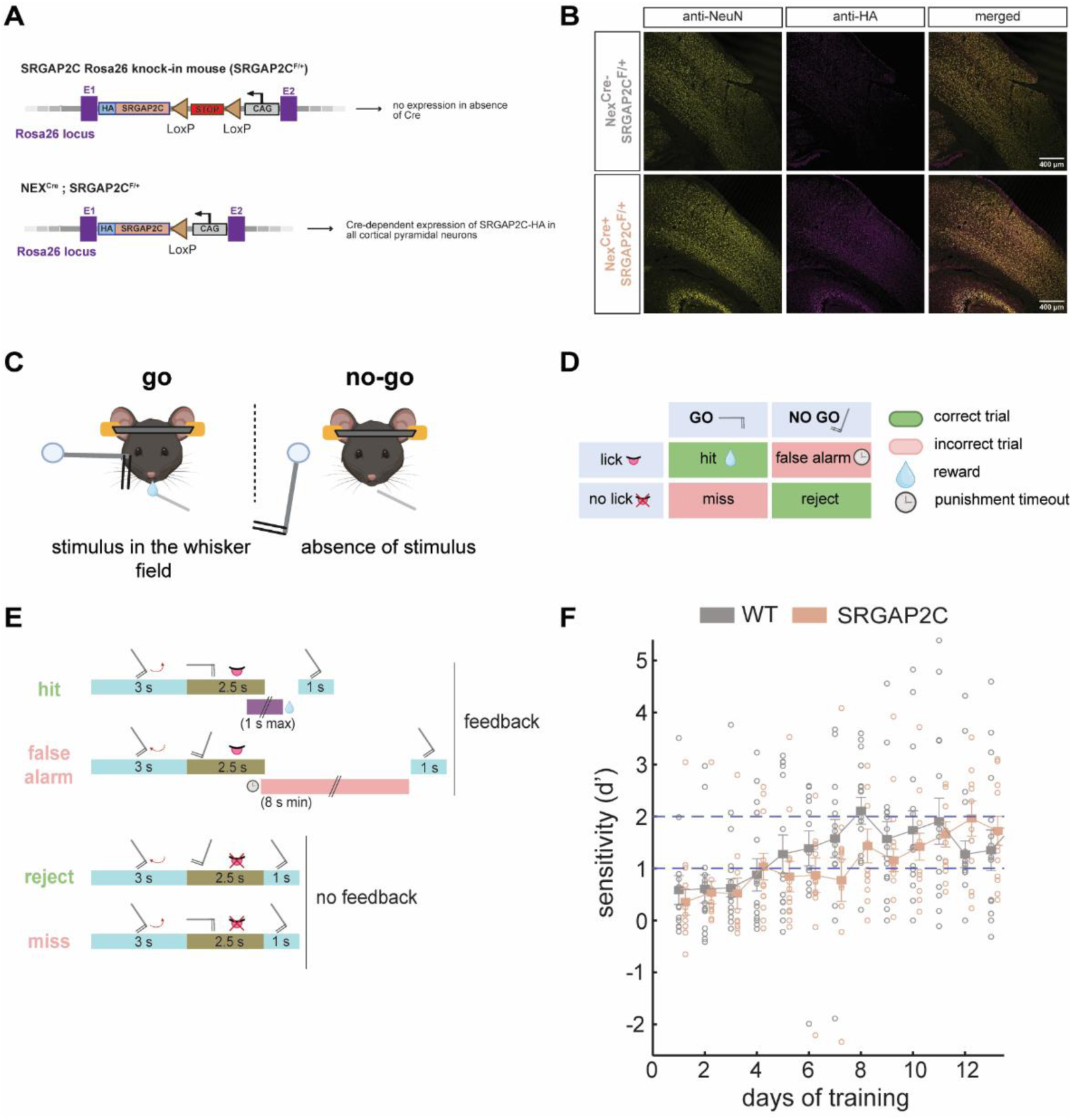
Experimental paradigm. A. Schematic representation of the transgenic knock-in strategy at the Rosa26 locus. SRGAP2C-HA is expressed in presence of Cre recombinase in all cortical pyramidal neurons. B. Immunostaining for HA on cortical coronal brain sections from P60 Nex^Cre+^ SRGAP2C^F/+^ and littermate control SRGAP2C^F/+^. NeuN (yellow) as a neuronal marker, HA (magenta) as reporter for SRGAP2C. C. Schematic representation of the behavioral sensory Go/NoGo task. Mice have to learn to lick when detecting the stimulus in their whisker field to receive a water reward, and refrain from licking in absence of the stimulus. The mice are head-fixed in a custom-built experimental set-up containing a small chamber, a water-port is positioned in front of their snout, and a metallic pole is moving on their right side. D. Possible outcomes for each trial type according to the mouse’s actions. Punishment timeout prevents the start of the next trial and is enacted until the mouse stops licking for 8 seconds. E. Trial structure in time. A hit provides feedback in the form of a water reward, a false alarm provides feedback by delaying the start of the next trial. Rejects and misses do not provide any feedback information. F. Sensitivity measured with the discriminability index (d’) across days of training. WT (n=14) and SRGAP2C mice (n=13) are shown in grey and light beige, respectively (genotype: F(1, 325) = 0.8, *p* = 0.37, day: F(12, 325) = 2.5, *p* < 0.005, genotype x day: F(12, 325) = 0.89, *p* = 0.56, linear mixed model and post-hoc ANOVA).

To evaluate the effect of human-specific SRGAP2C expression in cortical circuits on behavioral performance and learning strategy, we compared SRGAP2C mice with wild-type (WT; SRGAP2C ^F/+^ mice) in an active Go/NoGo sensory detection task. Water deprived mice of both genotypes have to actively detect the presence of a pole in using their whiskers and learn to associate this sensory cue with licking a port to receive a water reward or withhold licking in absence of the stimulus to avoid a punishment timeout (**Fig 1C-D**). Missed detections were not punished and correct rejections were not rewarded (**Fig 1D**). As a result, only the choice to lick provides any feedback in the form of a timeout and/or reward, while the choice not to lick has no instrumental effect on the trial (**Fig 1E**). Prior to beginning of training, mice were habituated to our custom experimental set-up and to water deprivation. To characterize the effects of SRGAP2C expression in cortical circuits on learning, we limited pre-training phases to a minimum and observed the strategy used to reach expert level behavior.

Based on prior work showing an increase in sensory coding by cortical L2/3 PNs in these SRGAP2C mice compared to WT control mice (*39*), we first determined whether SRGAP2C alters perceptual sensitivity. This was defined as the ability to discriminate between the presence and absence of stimulus quantified using the standard discriminability index (d’; see **Methods**) (*44*, *45*). We found that WT and SRGAP2C mice reach similar sensitivity levels across training, with both genotypes reaching thresholds of d’=1 (corresponding to about 69% correct responses) after 4 days (Fig 1E). Linear-mixed modelling showed no effect of genotype on sensitivity over time, with both groups showing similar improvement across days of training (genotype: F(1, 325) = 0.8, *p* = 0.37, day: F(12, 325) = 2.5, *p* < 0.005, genotype x day: F(12, 325) = 0.89, *p* = 0.56, linear mixed model and post-hoc ANOVA). This suggests that both groups learn to perform the task with similar sensitivity levels.

### Enhanced licking behavior in SRGAP2C mice

Despite having comparable sensory sensitivity (d’ scores), we did observe an increase in licking behavior in SRGAP2C mice, compared to WT (representative examples in **Fig 2A-B**). This implies that there may have been systematic differences in the strategy used by the two genotypes to achieve the same level of performance, therefore we next characterized their learning strategies. In general, behavior could be decomposed into two phases. First, we noticed an early learning phase in which both genotypes learned to lick more frequently in this task, as indicated by an increase in hit rate that happened in parallel with a decrease in the correct reject rate (**Fig 2C**). In the second learning phase, the hit rate remained stable and close to maximum for the rest of training while the correct reject rate slowly increased as the animals learned to selectively withhold licking in NoGo trials (**Fig 2C**).

**Figure 2.**
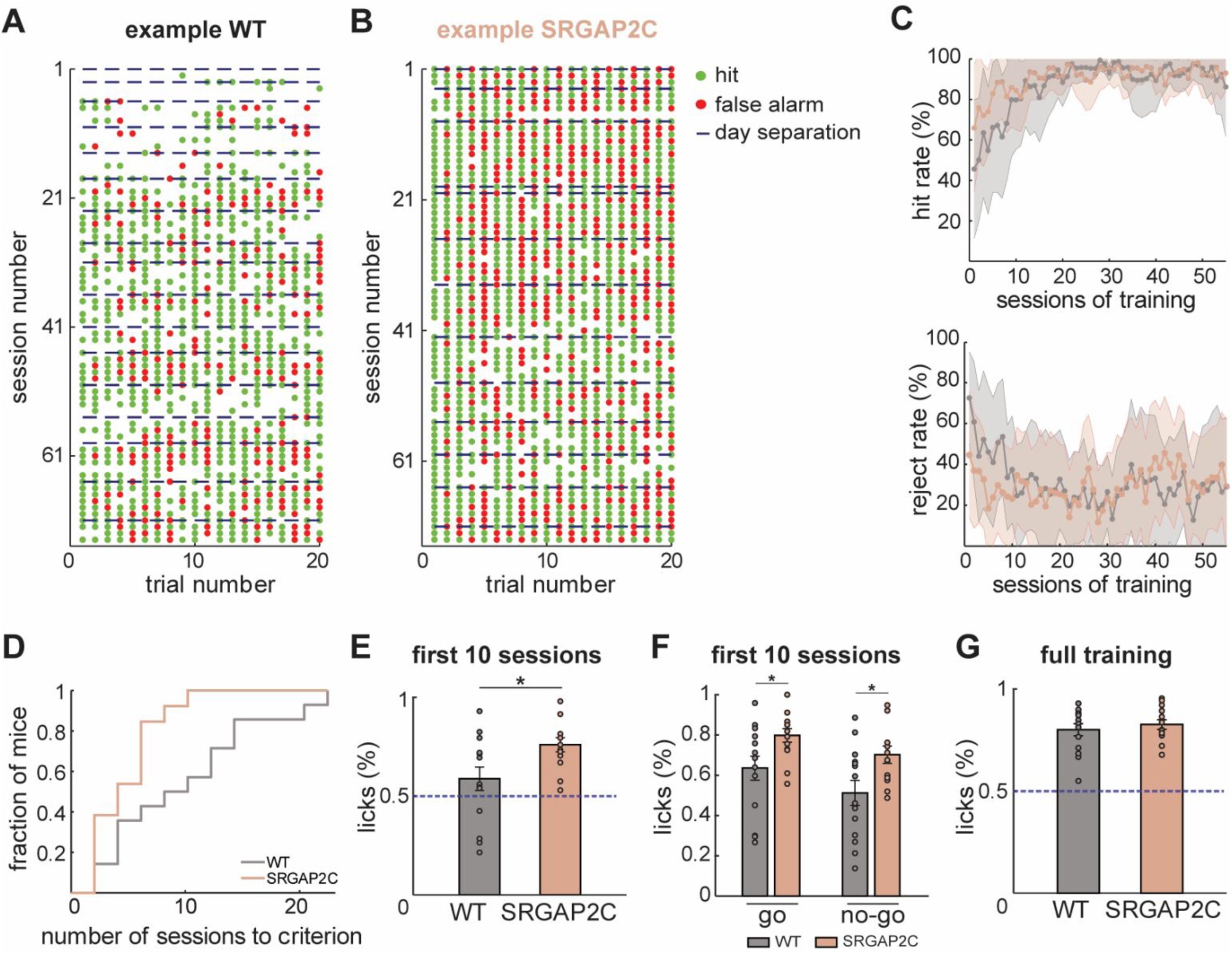
Enhanced licking behavior early in training for SRGAP2C mice. WT (n=14) and PNs-SRGAP2C (n=13) are shown in grey and pink beige respectively. A. Representative example of licking behavior for one example WT mouse. Raster plot of first licks showing each session of 20 trials on the y-axis and sessions in chronological order on the x-axis. Each black dot represents a lick. The same plot is shown on the right with hit trials in green and false alarms in red. Blue dotted lines represent day separations. B. Same for an example SRGAP2C mouse. C. Hit and reject rate across task learning. Hit rate in Go trials, calculated as the number of hits in presence of stimulus. Reject rate in NoGo trials, corresponding to the number of correct rejects in absence of stimulus. D. Cumulative stair plot of the number of sessions required to reach the criterion (first hit rate session above 90%). Significance was tested with a logrank test (mean WT: 8.36 ± 6.53 s.d, mean SRGAP2C: 3.39 ± 2.66 s.d, *p* < 0.05). E. Average proportion of licks for the first 10 sessions. Significance was tested with a Wilcoxon ranksum test (mean WT: 0.59 ± 0.22 s.d, mean SRGAP2C: 0.76 ± 0.13 s.d, *p* < 0.05, Wilcoxon ranksum test). F. Average proportion of licks for the first 10 sessions, separated by go and no-go trials. Significance was tested with a Wilcoxon ranksum test (go: mean WT: 0.64 ± 0.22 s.d, mean SRGAP2C: 0.8 ± 0.12 s.d, *p* < 0.05; no-go: mean WT: 0.51 ± 0.23 s.d, mean SRGAP2C: 0.7 ± 0.15 s.d, *p* < 0.05). G. Average proportion of licks across training. Significance was tested with Wilcoxon ranksum test (mean WT: 0.8 ± 0.11, mean SRGAP2C: 0.83 ± 0.08, *p* = 0.72, Wilcoxon ranksum test).

Although the same two learning phases were present for both genotypes, the hit rate for SRGAP2C mice started at significantly higher than WT (WT: 45.6 %, SRGAP2C: 65.87 %) and increased faster in the first 10 sessions (**Fig 2C**). This was very consistent across mice as a higher fraction of SRGAP2C mice reached a 90% hit rate criterion in very few sessions compared to controls (**Fig. 2D**; mean WT: 8.36 ± 6.53 s.d, mean SRGAP2C: 3.39 ± 2.66 s.d, *p* < 0.05, logrank test). This happened because SRGAP2C mice initiated licking in a higher proportion of trials in the first 10 sessions compared to WT mice (**Fig 2E**; mean WT: 0.59 ± 0.22 s.d, mean SRGAP2C: 0.76 ± 0.13 s.d, *p* < 0.05, Wilcoxon ranksum), with initiating lick proportion reaching above chance levels (50%) in all mice (**Fig 2E**). SRGAP2C mice licked more frequently independently of trial type, which suggests an enhanced drive to lick in both Go and NoGo trials (**Fig 2F**; go: mean WT: 0.64 ± 0.22 s.d, mean SRGAP2C: 0.8 ± 0.12 s.d, *p* < 0.05, Wilcoxon ranksum; no-go: mean WT: 0.51 ± 0.23 s.d, mean SRGAP2C: 0.7 ± 0.15 s.d, *p* < 0.05, Wilcoxon ranksum). This did not appear to be a generalized increase in licking in SRGAP2C mice because, across the whole of training, both groups initiated licks in similar proportions (**Fig 2G**; mean WT: 0.8 ± 0.11, mean SRGAP2C: 0.83 ± 0.08, *p* = 0.72, Wilcoxon ranksum). Instead, SRGAP2C mice seemed to reach and progress through the first learning phase faster than WT mice.

### Differences in licking are not driven by differences in thirst, compulsivity, or sensitivity to punishment

Although licking was useful in the task and could play an important part of how all mice acquired stimulus-action-outcome associations, it remained possible that SRGAP2C mice licked more initially because of low-level differences in thirst level, spontaneous repetitive behaviors, or insensitivity to punishment. This was not the case. First, there were no differences between genotypes in the effects of water deprivation. We measured the water intake 2 and 24 hours after the end of training (after the return to ad libitum water in the homecage) for both groups (Fig 3A). Genotype had no significant effect on water intake in either window (estimate = -0.81, 95% CI = [-3.12; 1.5], t (22) = -0.73, *p* = 0.47, linear mixed model; post-hoc comparisons 2h: *p* = 0.54; 24h: *p* = 0.69; after Bonferroni correction 2h: *p* = 1; 24h: *p* = 1). Thus, higher licking behavior in SRGAP2C mice compared to WT control mice is unlikely to be due to differences in satiation or thirst.

**Figure 3.**
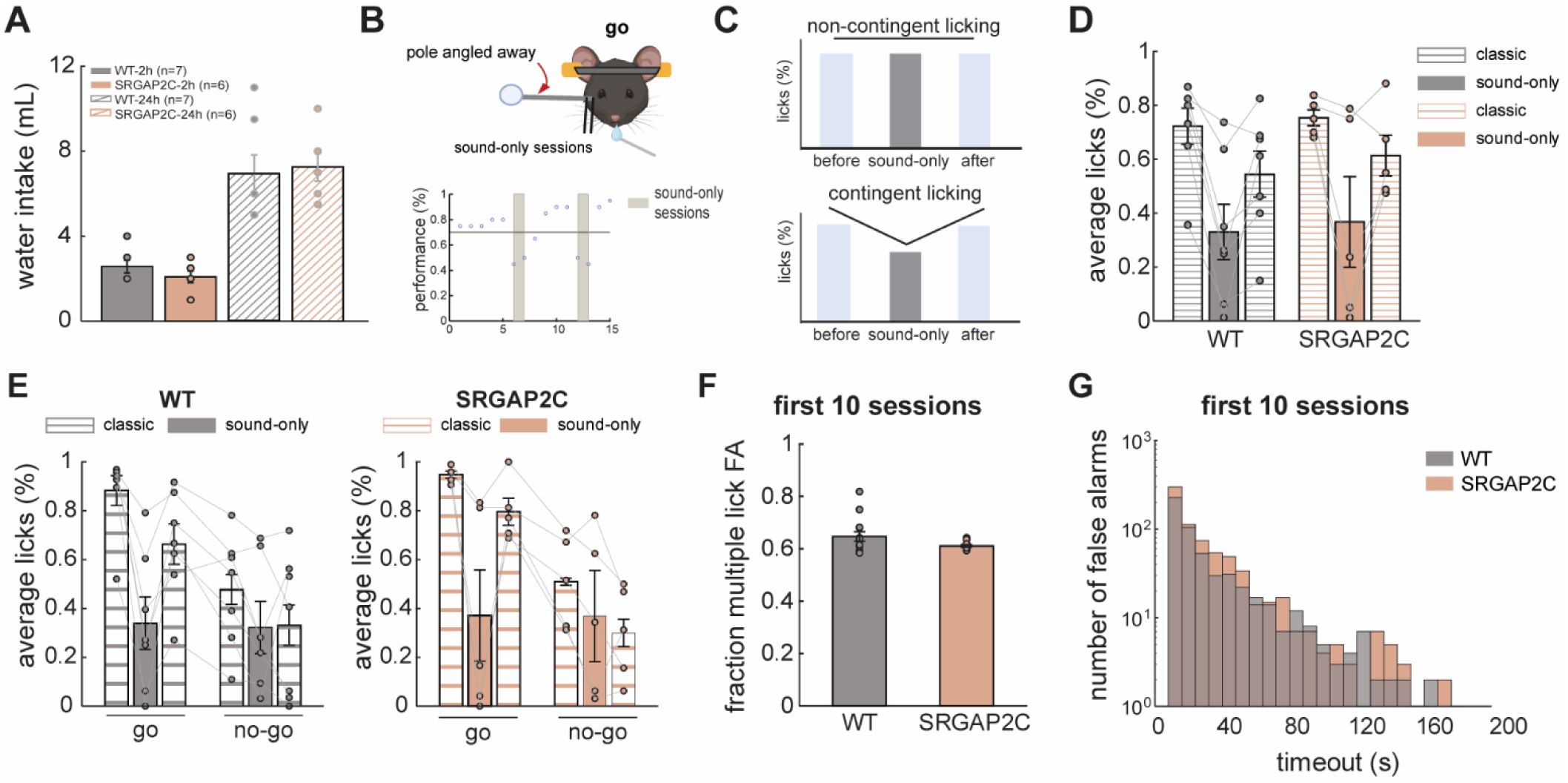
Licking is informative and whisker-dependent. A. Water intake after stop of water deprivation protocol. WT (n=7) and SRGAP2C (n=6) mice are shown in grey and pink beige respectively. A linear-mixed model with individual mice as grouping variable showed no significant change in water intake between genotypes (estimate = -0.81, 95% CI = [-3.12; 1.5], t (22) = -0.73, *p* = 0.47, linear mixed model). We compared water consumption between genotypes at the short (2h) and long (24h) recovery timepoints and confirmed that both groups are similarly affected (post-hoc comparisons, 2h: *p* = 0.54; 24h: *p* = 0.69; after Bonferroni correction 2h: *p* = 1; 24h: *p* = 1). B. *Top*. Schematic representation of the sound-only control experiment. The pole was angled away from the whiskers towards the experimenter, such that the mouse would not receive any somatosensory stimulus. *Bottom*. Typical performance profile for sound control experiment for a WT mouse. For 2 consecutive days, expert mice alternated between performing the classic task and sound-only sessions. Performance is plotted with blue dots as a function of session number and sound-only sessions are represented by the gray shaded areas. C. Schematic plot of licking contingency. Licking is a predictive cue if it decreases in sound-only sessions, also indicating whisker-dependency. D. Average proportion of licks in sound-only control experiment. For both genotypes, the bars represent the average lick proportion before, during and after the sound-only sessions. Before and after sessions (classic) are shown with bars filled with dashed horizontal lines and sound-only sessions are represented by filled color bars. WT (n=7) and SRGAP2C (n=5) mice are shown in grey and pink beige respectively. Linear mixed model and post-hoc ANOVA (session: F(2,30) = 10.31, *p* < 0.0005, genotype: F(1,30) = 0.06, *p* = 0.80, session x genotype: F(2,30) = 0.06, *p* = 0.94; post-hoc comparisons were collapsed by genotype, before/sound-only: estimate = -0.39, 95% CI = [-0.56; -0.21], t(30) = - 4.49, *p* < 0.0001; sound-only/after: estimate = -0.25, 95% CI = [-0.42; -0.07], t(30) = -2.86, *p* < 0.01, before/after: estimate = -0.14, 95% CI = [-0.32; 0.036], t(30) = -1.63, *p* = 0.11; after Bonferroni correction, before/sound-only: *p <* 0.0005; sound-only/after: *p* < 0.05, before/after: *p* = 0.23). E. Same as D but separating licks according to trial type. Linear mixed model and post-hoc ANOVA for hits (session: F (2,30) = 17.45, *p* < 0.0001, genotype: F (1,30) = 0.26, *p* = 0.61, session x genotype: F (2,30) = 0.3, *p* = 0.74) and false alarms (session: F (2,30) = 3.002, *p* = 0.065, genotype: F (1,30) = 0.054, *p* = 0.82, session x genotype: F (2,30) = 0.26, *p* = 0.77). F. Fraction of multiple lick false alarms (calculated by sessions) for the first 10 sessions of training. WT (n=14) and SRGAP2C (n=13) mice are shown in grey and pink beige respectively. Significance was tested with a Wilcoxon ranksum test: mean WT: 0.65 ± 0.067 s.d, mean SRGAP2C: 0.61 ± 0.019, *p* = 0.22. G. Distribution of false alarm durations in the first 10 sessions of training. Timeout duration refers to the time between first lick and end of trial. Y-axis shown in log scale. Bin size = 8 s. WT (n=14) and SRGAP2C (n=13) mice are shown in grey and pink beige respectively. Significance was tested with a K-S test: D = 0.04, n(WT) = 560, n(SRGAP2C) = 718, *p* = 0.69.

SRGAP2C mice also did not display a higher spontaneous licking behavior. A cohort of expert mice were presented with a variation of the task in which the whisker stimulus was removed, though they could still hear the movement of the apparatus (**Fig 3B**). If SRGAP2C mice simply tended to lick compulsively regardless of somatosensory cues, we would expect to see (1) no difference in licking behavior in sound-only sessions, compared to classic sessions before and after (**Fig 3B-C**), and (2) an increase in licking in SRGAP2C mice, compared to WT. Instead, both genotypes licked less and did so by a similar amount in sound-only sessions (**Fig 3D**). Linear mixed modeling revealed a significant effect of session identity (based on pole presence) on average lick rate but no difference based on genotype (session: F(2,30) = 10.31, *p* < 0.0005, genotype: F(1,30) = 0.06, *p* = 0.80, session x genotype: F(2,30) = 0.06, *p* = 0.94, linear mixed model and post-hoc ANOVA; post-hoc comparisons were collapsed by genotype, before/sound-only: estimate = -0.39, 95% CI = [-0.56; -0.21], t(30) = - 4.49, *p* < 0.0001; sound-only/after: estimate = - 0.25, 95% CI = [-0.42; -0.07], t(30) = -2.86, *p* < 0.01, before/after: estimate = -0.14, 95% CI = [-0.32; 0.036], t(30) = -1.63, *p* = 0.11; after Bonferroni correction, before/sound-only: *p <* 0.0005; sound-only/after: *p* < 0.05, before/after: *p* = 0.23). Separating this data between Go and NoGo (sound-only) trials also did not reveal any differences by genotype (**Fig 3E**, for hits, session: F(2,30) = 17.45, *p* < 0.0001, genotype: F(1,30) = 0.26, *p* = 0.61, session x genotype: F(2,30) = 0.3, *p* = 0.74) and false alarms (session: F(2,30) = 3.002, *p* = 0.065, genotype: F(1,30) = 0.054, *p* = 0.82, session x genotype: F(2,30) = 0.26, *p* = 0.77, linear mixed model and post-hoc ANOVA). Together, these results suggest that SRGAP2C mice do not display increased compulsion to lick but instead, they licked selectively when licking could elicit information about the state of the world.

Finally, we considered the possibility that SRGAP2C mice licked more because they were insensitive to the timeout punishments. While licking in a NoGo trial always resulted in a timeout punishment, the duration of punishments varied, depending on subsequent licks during timeout. Mice could either lick exactly once and be minimally punished with an 8 second timeout, or they could continue to lick repeatedly, which would increase the timeout duration until the point at which they were able to successfully withhold licking for 8 consecutive seconds. If licking behavior of SRGAP2C mice was compulsive or driven by an insensitivity to punishment, we would expect these mice to continue licking during the timeout and to experience longer punishment durations as a result. Instead, in the first 10 sessions, when the SRGAP2C animals licked the most, the proportion of false alarm trials with multiple licks was not significantly different for the two genotypes (**Fig. 3F**, mean WT: 0.65 ± 0.067 s.d, mean SRGAP2C: 0.61 ± 0.019, *p* = 0.22, Wilcoxon ranksum). In addition, both groups spent a similar amount of time in timeout, as timeout durations distributions are statistically comparable (**Fig 3G**, D = 0.04, n(WT) = 560, n(SRGAP2C) = 718, *p* = 0.69, K-S test). Again, these results suggest that the increased licking in the SRGAP2C mice had an instrumental function: it only happened early in learning, was not connected with differences in thirst, but instead was selective to the specific epoch of the task in which it was most useful.

### SRGAP2C promotes information accumulation

The SRGAP2C mice used a strategy that initially led to more punishments but did not seem otherwise less sensitive to punishment. One possibility is that the SRGAP2C mice used this strategy because licking constitutes the most informative gathering action in this task, particularly before the acquisition of stimulus-action-outcome associations. Indeed, when we calculated the Shannon information (*46*) of the mice’s action sequences (**Fig. 4A**; see **Methods**), we found that SRGAP2C mice accumulated a larger amount of information compared to WT mice. In fact, cumulative information gain of WT mice was comparable to that of a simulated random strategy (**Fig 4A**). Furthermore, while cumulative information predicted task performance in WT mice, it remains high and stable in SRGAP2C mice (**Fig 4B**; WT: r = 0.71, *p* < 0.0001, SRGAP2C: r = 0.21, *p* < 0.05, linear regression). These results suggest that early licking in SRGAP2C mice could reflect an information-seeking strategy.

**Figure 4.**
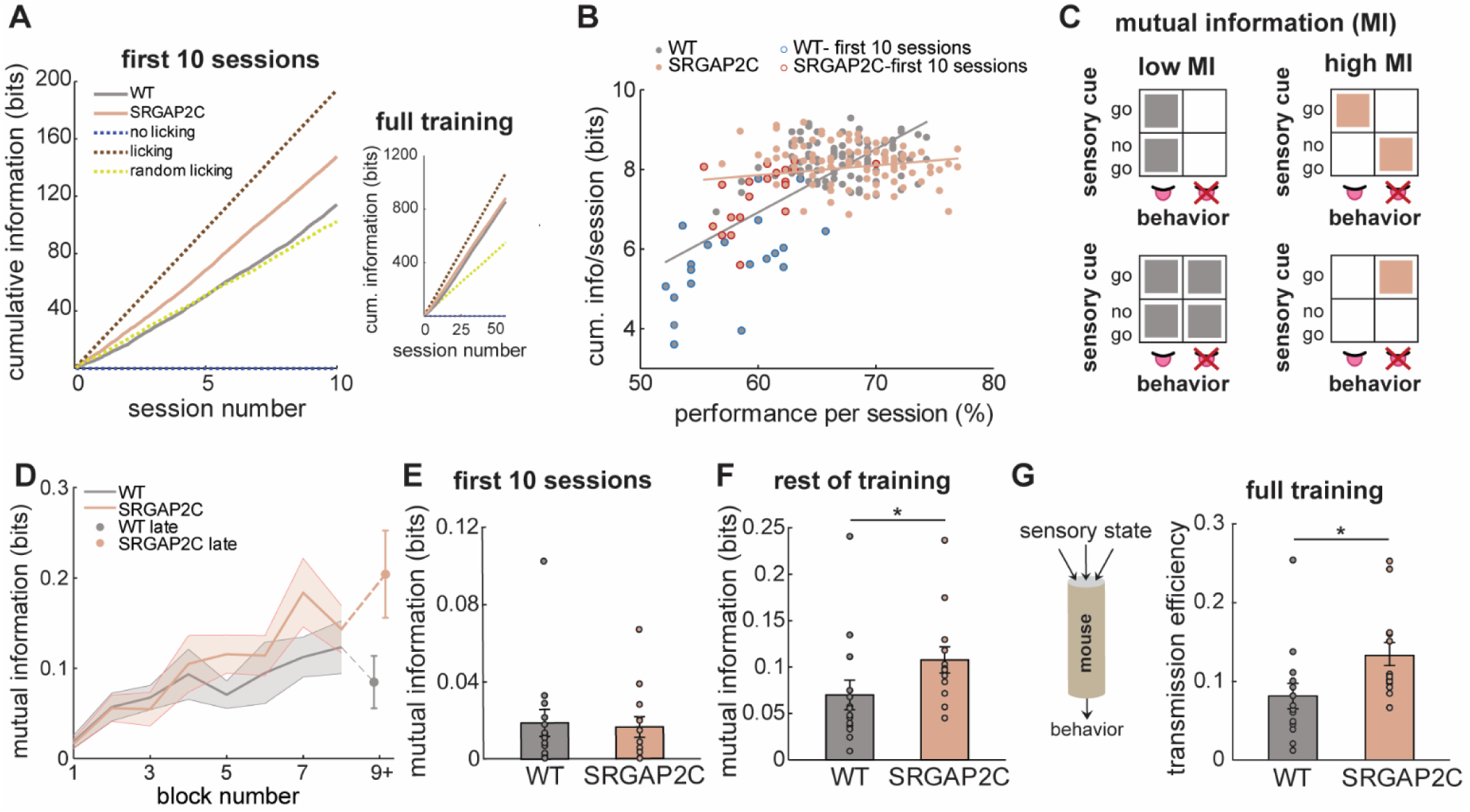
SRGAP2C mice favor an information-seeking strategy. A. Amount of information accumulated over time during the first 10 sessions, inset shows the whole training period. WT (n=14) and SRGAP2C (n=13) are shown in grey and pink beige respectively. Dotted lines indicate simulations for no licking, licking 100% of the time and random licking in blue, brown and yellow, respectively. B. Correlation between performance and cumulative information. WT (n=14) and SRGAP2C (n=13) are shown in grey and pink beige respectively. Linear regression: WT: r = 0.71 (*p* < 0.0001), SRGAP2C: r = 0.21 (*p* < 0.05). C. Schematic representation of low and high mutual information (MI) in a Go/NoGo task. Low MI indicates a behavior unrelated to the input (always licking or never licking independently of trial type), whereas high MI indicates a discriminative behavior (licking or not licking is associated with trial type). D. Mutual information across training. WT (n=14) and SRGAP2C (n=13) are shown in grey and pink beige respectively. Trials were grouped in blocks of 10 sessions (200 trials). For the late phase (blocks ≥9), all trials were pooled across mice due to low per-block sample size. Linear mixed model: genotype: *p* = 0.36, block: *p* < 0.0001, genotype x block: *p* < 0.005. Significance on the last block was tested with a t-test: mean WT (n=5): 0.08 ± 0.03 s.d, mean SRGAP2C (n=6): 0.2 ± 0.05 s.d, *p* = 0.075. E. Mutual information during the first 10 sessions. WT (n=14) and SRGAP2C (n=13) are shown in grey and pink beige respectively. Significance was tested with a Wilcoxon ranksum test: mean WT: 0.019 ± 0.026 s.d, mean SRGAP2C: 0.016 ± 0.019 s.d, *p* = 0.75. F. Same as E but for the rest of the training period. Significance was tested with a Wilcoxon ranksum test: mean WT: 0.07 ± 0.059 s.d, mean SRGAP2C: 0.11 ± 0.05 s.d, *p* < 0.05. G. *Left*. Schematic representation of an information channel. *Right.* Transmission efficiency (defined as mutual information divided by output entropy) across training. WT (n=14) and SRGAP2C (n=13) are shown in grey and pink beige respectively. Significance was tested with a Wilcoxon ranksum test: mean WT: 0.08 ± 0.06 s.d, mean SRGAP2C: 0.13 ± 0.06 s.d, *p* < 0.05.

### SRGAP2C facilitates preservation of task-relevant information

Although discriminability did not differ by genotype in expert mice (**Fig 1E**), it remained possible that differences in strategy persisted throughout the task. To test for this possibility, we examined the mutual information between each action (licking or not licking) and its perception (stimulus presence or absence). The discriminability index is a measure of correct performance, whereas mutual information measures the extent to which behavior is structured by sensory information.

Because mutual information increases when there is a strong relationship between the licking behavior and perception (**Fig. 4C**), we first estimated the mutual information between sensory input (trial type) and motor output (mouse’s actions) across blocks of 200 trials (10 sessions). As expected, we found that mutual information increased over time with learning in both genotypes (**Fig 4D**; block: f (1,220) =93.44, *p* < 0.0001; linear mixed model and post-hoc ANOVA). Further, while there was no significant difference between the genotypes, mutual information increased more rapidly in the SRGAP2C mice than in the WT mice (genotype: f (1,220) = 0.85, *p* = 0.36, genotype x block: f (1,220) = 9.98, *p* < 0.005). SRGAP2C mice also showed a trend for higher mutual information even in very late stages of learning that were only measured in a subset of animals (**Fig 4D**; late combined block: mean WT (n=5): 0.08 ± 0.03 s.d, mean SRGAP2C (n=6): 0.2 ± 0.05 s.d, *p* = 0.075, t-test). These effects were driven by long-term differences in task strategy, not differences in early learning. No genotype differences were observed in the first 10 sessions (**Fig 4E**; mean WT: 0.019 ± 0.026 s.d, mean SRGAP2C: 0.016 ± 0.019 s.d, *p* = 0.75, Wilcoxon ranksum), but SRGAP2C mice showed significantly higher mutual information during the rest of training compared to WT (**Fig 4F**; mean WT: 0.07 ± 0.059 s.d, mean SRGAP2C: 0.11 ± 0.05 s.d, *p* < 0.05, Wilcoxon ranksum; no significant genotype effect in shuffled data: **Supp Fig 1**). This suggests that the behavior of SRGAP2C mice becomes more tightly organized with respect to the stimulus across learning compared to WT, which might in part explain a trend for higher sensitivity levels after weeks off task (**Supp Fig 2**).

Although prior work reported an increased reliability of sensory coding of L2/3 PNs in SRGAP2C mice compared to WT control mice (*39*), we observed no improvement in sensory sensitivity measured behaviorally. The increase in mutual information characterizing the behavioral strategy used by SRGAP2C mice did raise an intriguing alternative view, however: perhaps the changes in cortical circuit connectivity characterizing the SRGAP2C mice globally improve the neural signal-to-noise ratio rather than specifically improving sensory coding. If this were the case, we would expect these mice to be better information channels, not simply more perceptually accurate. To test this idea, we calculated the transmission efficiency of SRGAP2C and WT mice: a measure of how efficiently information is transformed from input to output across a communication channel (see **Methods**). We found that transmission efficiency was significantly higher in SRGAP2C mice compared to WT across learning (**Fig 4G**; mean WT: 0.08 ± 0.06 s.d, mean SRGAP2C: 0.13 ± 0.06 s.d, *p* < 0.05, Wilcoxon ranksum). Together, these results suggest that SRGAP2C mice not only favor an information-seeking strategy, but are themselves better information-preserving channels, consistent with prior observations that expression of the SRGAP2C in cortical pyramidal neurons increases signal-to-noise ratio of sensory-evoked response i.e. cortical neurons in SRGAP2C mice are more accurate at coding the presence or absence of sensory cues (*39*).

### Modeling cognitive flexibility replicates SRGAP2C mice’s behavior

SRGAP2C mice both sought more information, better preserved information, and learned faster compared to WT mice. In order to determine whether they could be related in some way, we turned to reinforcement learning (RL). Specifically, we considered a framework in which agents learn to generate reinforced responses in the face of limitations on the complexity of their policies (see **Methods**) (*47*). In the policy compression framework, a policy—an agent’s mapping between states and actions—can be understood, abstractly, as a communication channel that transforms information about the state of the world (the sensory input) into an action (the motor output). Policy compression defines the complexity of the policy as the capacity of the information channel: its ability to preserve the information that passes through it without loss (i.e. without compression). If an agent has a low capacity (high policy compression), its ability to flexibly guide action in response to a specific state is limited and the animal tends to do the same thing, regardless of circumstance (**Fig 5A**). Action selection is a habitual process that is virtually independent from the state: an outside observer would not be able to guess the state identity by observing the action. On the other hand, if an agent has a high capacity (low policy compression), it would be able to more easily depart from habitual actions because it can represent the more complex set of associations between states and actions needed to perform a state-specific action.

**Figure 5.**
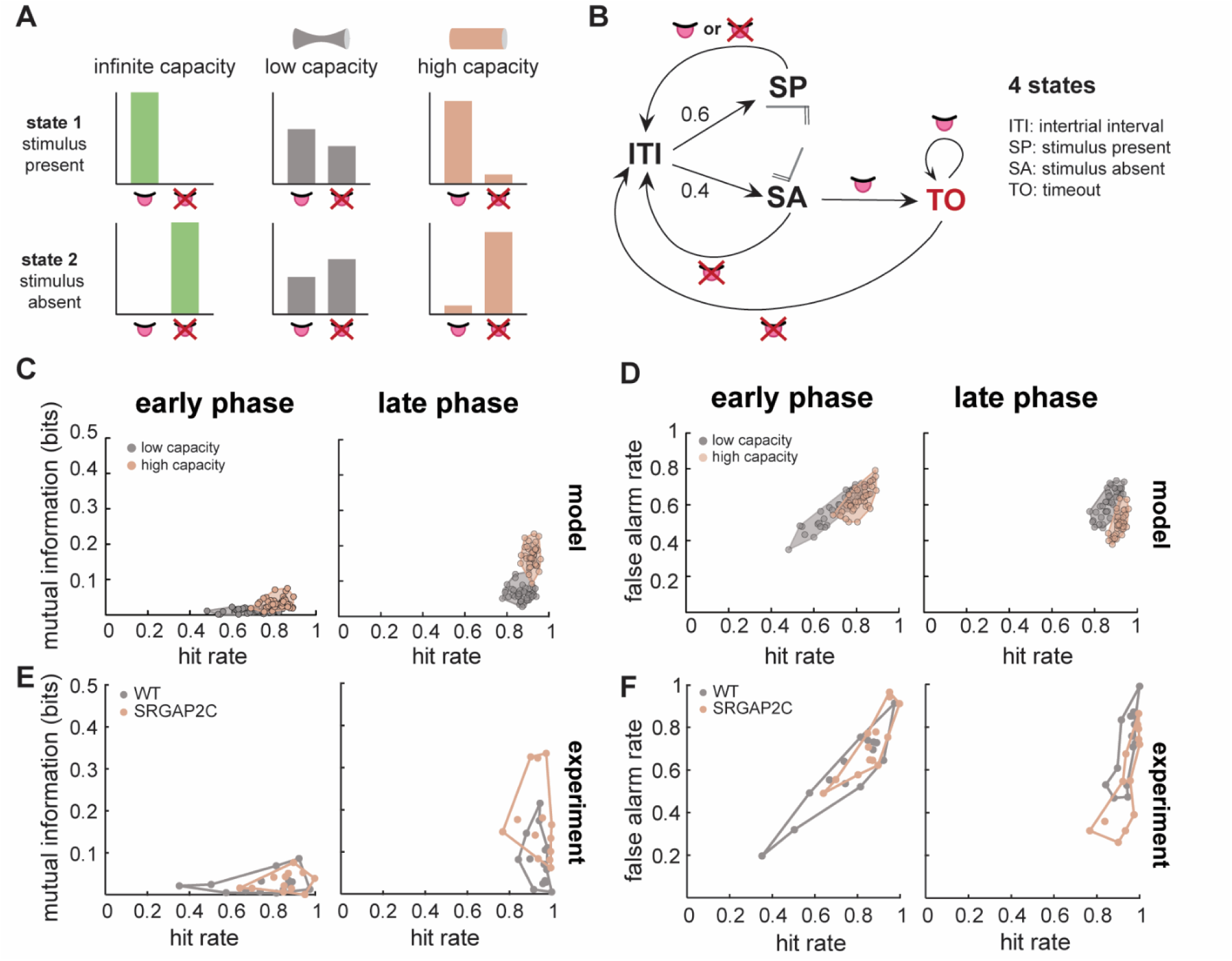
Key features of SRGAP2C mice behavior are replicated by a reinforcement learning model. A. Schematic representation of the notion of capacity in the model: a low-capacity agent will tend to have habitual actions regardless of state identity (high policy compression). A high-capacity agent will take state-specific actions (low policy compression). B. Schematic representation of the model task epochs and their transitions. C. Contour plots of mutual information in function of hit rate comparing early and late training phases, obtained in RL model simulations. D. Contour plots of false alarm rate in function of hit rate comparing early and late training phases in RL model simulations. E. Corresponding contour plots of C obtained out of experimental data. F. Corresponding contour plots of D obtained out of experimental data.

We modeled the Go/NoGo task using four different states that correspond to the different task epochs (**Fig 5B**) and then trained both high- and low-capacity agents to perform the task. Note that all other parameters of the models, including learning rates, were identical between the two agent types—the only difference in the two agents was the extent of the limitation in the complexity of their policy (see **Methods**). As the agents learned, both high- and low-capacity agents progressed from an early phase of learning when hit rates were low towards a late stage of learning where hit rates were high (**Fig. 5C-D**). However, the high-capacity agents learned faster: their hit rates led the low-capacity agents’ hit rates during the early phase of learning before both types ultimately reached more comparable hit rates in the late phase of learning (**Fig. 5C**, x-axis). This parallels our observations that SRGAP2C mice learned to lick faster than WT mice (**Fig. 5E**; **Fig. 2C**, **Fig. 2E**) Because policy compression limits the mutual information between states and behavior by design, it was expected that high-capacity agents would have higher mutual information than low-capacity agents. However, this difference was only pronounced late in learning: mutual information was similar between the two types of agents early in learning (**Fig. 5C**). This parallels the fact that mutual information was only significantly elevated in SRGAP2C mice compared to WT late in learning, with few differences in early learning (**Fig. 5E**; **Fig. 4E-F**). Together, these results suggest that simply increasing the policy complexity of an RL agent is sufficient to recapitulate genotypic differences in the speed of learning and the emergence of structured behavior in SRGAP2C mice.

We next considered how the agents would differ in the reduction in false alarm rate that defined the transition from the early to late stage of learning in this task. Recall that early in learning, both genotypes move from a phase of indiscriminate licking into a phase in which they lick only in the presence of the stimulus. Both the high- and low- capacity agents were able to replicate this phenomenon (**Fig. 5D**), suggesting that a naïve RL model will adopt the same 2-phase strategy that the mice did in this task. However, there were also differences due to the capacity of the agents. In the early phase of learning, high-capacity agents displayed both a high hit rate and a high false alarm rate compared to the low-capacity agents (**Fig 5D**). In this task, this means high-capacity agents would display a high level of licking behavior. This paralleled the increase in licking we initially observed during early learning in SRGAP2C mice compared to WT (**Fig. 5F**; **Fig. 2C**, **Fig. 2E**). While the hit rate is high for both agent types in the late phase of learning, the high-capacity agents had a larger reduction in the false alarm rate, compared to the low-capacity agents. We found that SRGAP2C mice similarly had a lower false alarm rate than WT mice in the late phase of learning (**Fig. 5F**). Thus, manipulating only the capacity of otherwise identical RL agents reproduced every major difference between SRGAP2C and WT mice: from their early differences in indiscriminate licking to their late differences in the extent to which their behavior preserved the structure of the world.

## DISCUSSION

We evaluated the learning behavior of mice expressing the human gene *SRGAP2C* in cortical pyramidal neurons in an active Go/NoGo detection task. Although WT and SRGAP2C mice had similar perceptual sensitivity, SRGAP2C mice exhibited a large amount of licking early in training, which could not be explained by differences in motivation or compulsivity. The SRGAP2C mice did not lick excessively when doing so offered no informational advantage. While both SRGAP2C and WT mice eventually learned to withhold licks to avoid punishment, the behavior of SRGAP2C mice was more organized with respect to the environment throughout learning. Importantly, we were able to replicate all of these results with a simple reinforcement learning model in which SRGAP2C expression permits a more complex and lossless representation of the structure of the world. Together, these results suggest that human-specific SRGAP2C may play a role in increasing the capacity of cognition, the reliability with which behavior can be structured with respect to the environment.

Rather than simply improving task performance, SRGAP2C expression in cortical pyramidal neurons modified the strategy by which mice learned the task contingencies. While indiscriminate or seemingly compulsive behavioral strategies have traditionally been ascribed to inattention, motor errors or sensory defects, a growing body of evidence suggests that rodents use multiple goal-directed strategies when faced with uncertainty, balancing the maximization of known rewards and the exploration of new options to build an accurate model of the environment (*5*, *6*, *42*, *48*–*50*). Quantitative, information-theoretic analyses of the structure of this task revealed that enhanced licking in SRGAP2C mice in the early phases of learning allowed these mice to accumulate more information than WT mice. This suggests that the high false alarm rate observed in SRGAP2C mice may be better interpreted as an index of engagement or information seeking, rather than as a failure to understand task rules. Indeed, even naïve RL agents went through a phase of high licking when learning this task, although this was never expressly engineered into these agents. Ultimately, these findings are consistent with observations that humans and animals use strategies to acquire task rules that may be silent with respect to their motoric choices, which could be a form of latent knowledge acquisition (*51*, *52*) or a dimensionality reduction strategy that speeds learning through reducing the variables to which credit might be assigned (*42*). By actively and indiscriminately licking early in the task, SRGAP2C mice may have improved their ability to gain certainty about task parameters.

The information-seeking and information-preserving qualities of SRGAP2C mice raise the possibility that they have a general drive to gain information. In a word, these results could suggest that SRGAP2C mice are more curious than their WT counterparts. While there is no generally agreed upon definition of curiosity (*53*), the behavior of SRGAP2C mice does have some parallels with commonly identified features of curiosity. For one, the SRGAP2C mice performed informative actions despite costly punishments and a reduced rate of reward. This is consistent with observations from the curiosity literature that many animals will sacrifice positive outcomes in favor of novel information, including mice (*54*), rats (*55*), pigeons (*56*), and monkeys (*57–61*). However, curiosity is also thought to have an instrumental purpose: it should lead to better learning or improved long-term recall (*62*). Indeed, naive SRGAP2C mice learned faster, expert SRGAP2C mice showed an improved encoding of task structure, and we also found a tendency for enhanced long-term recall in SRGAP2C mice after weeks off task. Of course, SRGAP2C mice would need to be compared with WT in validated curiosity tasks to justify any speculation that SRGAP2C expression may play a role in human curiosity, but these results are not inconsistent with the idea that humans may have some specialization for curiosity (*63–65*), even as curiosity-like behaviors are certainly present in other species.

It is worth pondering how our observations in SRGAP2C mice may provide insights into human-specific cognitive evolution. In this respect, two observations stand out: (1) the fact that SRGAP2C mice were better information channels throughout training, and (2) that we were able to reproduce essentially all of their behavioral differences in RL agents in whom we only increased representational capacity. Taken together, these results are highly consistent, suggesting that the primary difference between SRGAP2C mice and their WT counterparts lies in their capacity to encode a richer representation of the states of the world. Of course, neither our mice, nor humans (*66*) represent the world in its full complexity, likely due to a conserved drive to balance maximizing reward with limiting cognitive load (*47*, *67–69*). Nonetheless, if one of the cognitive consequences of SRGAP2C expression is to increase the capacity of a brain to faithfully represent the world, it would resonate with observations in comparative cognition. We know, for example, that humans have an unusual capacity to share mental states and transmit information between themselves (*7–9*). Perhaps human social communication co-evolved with an SRGAP2C-dependent capacity to more accurately represent the world in our species—allowing human brains to rely on each other’s observations and beliefs more as our fidelity as information channels increased over evolutionary time.

The fact that SRGAP2C mice were better information channels in a sensorimotor learning task could be related to changes in sensory discrimination. Although we observed no genotype differences in the standard sensory discrimination index (d’), representing the state of the world in a perceptual task does require good sensory discrimination. Previous reports have shown that SRGAP2C expression increases response probability to passive whisker stimulation in layer 2/3 cortical pyramidal neurons in primary somatosensory cortex, which suggests an enhanced ability to encode sensory evidence (*39*). Highly-tuned processing in primary sensory areas could be sufficient to allow efficient encoding of evidence accumulation in higher-order areas like prefrontal and parietal cortices, which have been shown to be essential to perceptual learning and decision-making (*70–73*). In this view, the availability of early and accurate sensory information could facilitate better internal representations of the task that could in turn explain how early information-seeking becomes paired with structured use of task-relevant information in expert SRGAP2C mice. Alternatively, SRGAP2C could broadly increase the signal-to-noise ratio across brain regions responsible for learning and decision-making, increasing representational capacity via reducing the noise in sensorimotor transformations. Recent evidence in this sense was found through mesoscale imaging in this mouse line during a sensory discrimination paradigm, where SRGAP2C expression in cortical pyramidal neurons induces a redistribution of cortical network activity towards more associative areas during motor planning while also increasing the discriminability of representational brain states (*41*). Future work will need to determine the neural mechanisms for the effects we report here, but also whether similar differences in behavior are observed in SRGAP2C mice in tasks that capture the complexity of natural decision-making, via requiring that mice integrate cues from multiple modalities, deal with changing reward structures, interact with conspecifics, and actively avoid obstacles and predators.

Overall, this study shows that the expression of the human-specific gene SRGAP2C in cortical pyramidal neurons—which increased cortico-cortical connectivity—can modify the strategy by which mice learn and execute a sensorimotor task. In doing so, our results elaborate on recent work showing that the loss of the ancestral SRGAP2A—which phenocopies the effects of human-specific SRGAP2C expression (*34*)— enhances structural synaptic plasticity in layer 2/3 pyramidal neurons (*74*), while studies directly introducing SRGAP2C have demonstrated increases in sensory coding of L2/3 PNs (*39*), and large-scale reorganization of cortical response dynamics (*39*, *41*). SRGAP2C is only one of many human-specific genetic modifiers that appeared during human brain evolution. Its contribution in the learning and cognitive abilities of the human lineage has to be understood in concert with other genetic mechanisms that have modified features of the human brain, including increased production of L2/3 PNs, increased cortical connectivity, changes in pyramidal functional properties (*18*, *75*). Understanding the cognitive impacts of these modifications could someday help us understand human-specific learning abilities and developmental trajectories, linking these important specializations to their species-specific genetic and cellular determinants.

## METHODS

### Mice

All experimental procedures were performed in compliance with experimental protocols 2020-2036 and 2024-6788 approved by the *Comité Institutionnel de Bonnes Pratiques Animales en Recherche* (CIBPAR) at the Centre de Recherche Azrieli, CHU Sainte-Justine (QC, Canada) in accordance with the Canadian Council of Animal Care (CCAC).Transgenic knock-in Nex^Cre+^SRGAP2C ^F/+^mice are heterozygous for the transgene, which is expressed in all cortical and hippocampal pyramidal neurons (*39*, *76*). Control WT mice were littermates and include the Nex^Cre-^SRGAP2C^+/+^, Nex^Cre-^SRGAP2C^F/+^ and Nex^Cre+^SRGAP2C^+/+^ genotypes. Transgenic mice and their control littermates were housed on a 12-h light/dark cycle with ambient temperature 20 to 24 °C and 40 to 70% humidity. Male and female mice were used in comparable proportions.

### Immunostaining

#### Intracardiac perfusion and section preparation

Adult mice (P60-64) were anaesthetized with 3-4% isoflurane. An incision was made to expose the heart, and a small cut was made in the right atrium. 1X PBS (37°C) was injected in the left ventricle of the beating heart until organs appeared white, followed by injection of 4% paraformaldehyde (PFA) (4°C). The brain was then dissected and maintained in 4% cold PFA for 2 hours post-fixation. It was then washed 6 times in 1X PBS (30 minutes at 4°C) under gentle agitation, and finally dehydrated overnight in 15%, then 30% sucrose. Brain coronal sections (40 um) were prepared with a cryostat (CryoStar NX50, ThermoScientific).

#### Immunostaining and imaging

Tissue sections were thawed and hydrated in 1X PBS for 10 minutes. Sections were incubated in a blocking buffer (0.2% Tween20) for 1 hour. Next, primary antibodies were incubated overnight at 4°C in a humidified chamber. The following day, sections were washed 3 times for 10 minutes in 1X PBS, and subsequently incubated with secondary antibodies for 1 hour in a humidified chamber. Sections were washed in PBS 2 times for 10 minutes, then incubated with DAPI (1:1000, BioRad) for 5 minutes, after which sections were washed in PBS for 10 minutes. Slides were carefully dried and mounted with Immu-mount (Epredia). Antibodies used were mouse anti-HA (1:1000, MMS-101P Biolegend), rabbit anti-NeuN (1:500, Abcam ab104225), donkey anti-mouse Alexa Fluor 647 (1:200; 715605150 Jackson ImmunoResearch) and anti-rabbit Alexa Fluor 568 (1:200, 709296149 Jackson ImmunoResearch). Images were taken with a confocal microscope at 10x magnification (Leica Stellaris WLL+).

### Surgery for fixation bar implantation

Prior to surgery (1-2h), 2-4 months old mice were intraperitoneally injected with slow release buprenorphine (1 mg/kg). Anaesthesia was induced with 3-4% isoflurane with 3% O_2_ during 5 minutes and maintained at 1-2% isoflurane throughout surgery, reflexes being monitored regularly. Mice were placed on a heating pad for the duration of the procedure and their head was fixed in a stereotaxic apparatus. Enrofloxacin (5 mg/kg) was injected subcutaneously, after which the hairs above the head were removed and betadine was applied. Bupivacaine (2.5 mg/kg) was injected subcutaneously at the top of the skull, and an incision was made to expose the dorsal skull. 20 minutes after the incision, carprofen (5 mg/kg) was injected subcutaneously. Conjunctive tissue was removed with tweezers and scissors. Once the skull surface had dried, it was hatched with a scalpel and veterinary tissue adhesive (3M Vetbond 1469SB) was applied. Posteriorly, in front of the ears, superglue (Lepage 2598431) was applied, after which the fixation bar was placed on top. Acrylic denture was used to seal the whole area. Mice were allowed to recover in a warm cage until fully awake, they were then transferred to individual housing and given 7 days to recover before the habituation protocol began. During this time, mice were monitored for any signs of stress, unease or infection at the operated site. Their weight was monitored to ensure a weight loss inferior to 15% of post-surgery weight.

### Water deprivation and habituation

Water was given every day in a petri dish, decreasing the amount of 0.5 mL every two days from the initial 2.5 mL until reaching the minimal volume of 1 mL. This corresponds to 20 to 35 % of their water consumption ad libitum (*77*). Mice were handled by the experimenter every day for 5-10 mn until they were able to groom in the experimenter’s hand. They were head-fixed progressively, slowly increasing the amount of time spent in the apparatus.

### Behavioral setup

The behavioral setup is a custom-made system. Mice are head-fixed in a small chamber, and a water delivering tube or port is placed in front of their mouth. A solenoid valve (Lee Company, LHLA1231111H) connected to the water port triggers the delivery of the water reward. A metallic pole composed of two vertical pipes of diameter 1.5 mm is positioned on the right side of the mouse, and moves thanks to a stepper motor (1205-POLOLU from Abra Electronics). Each component is controlled via an Arduino Uno and OM2 boards and custom-written Arduino and Matlab codes.

### Go/NoGo task

#### Pole movement

Mice had to detect the presence of a pole moving in and out of their whisker field. The pole has a resting position at a clockwise 60° angle from the plane. In the Go position, the pole moves counter-clockwise by 30° until it reaches a position anterior and lateral with respect to the snout, so that the mouse has to whisk actively in order to detect the pole. The calibration was specific to each mouse and adjusted prior to each session. A lick in the Go positioned triggered reward delivery. In the NoGo position, the pole moved clockwise by 30° in a position where the mice could not detect the pole. The task was performed in the dark, in order to ensure no visual cue could predict the movement of the pole prior to it reaching the whisker field. A lick in the NoGo position triggered a punishment timeout, mice had to withhold licking for 8 seconds to enable the initiation of the following trial. Trial type presentation was random, with 60% of Go (Real) trials and 40% of NoGo (Sham) trials. Reward was randomly delayed to 0.25, 0.5, 0.75 or 1 s after a correct lick.

#### Training

During the first three days of training, mice experienced “help” sessions in which the reward was always delivered in presence of the stimulus regardless of the mice’s licking behavior (day 1: 3 help sessions, day 2: 2 help 1 regular sessions, day 3: 1 help 2 regular sessions). Starting from day 4, mice were presented with regular sessions only. Help sessions were excluded from the analysis, but all regular sessions were included. Each day, mice were presented with an average of 5 regular sessions of 20 trials (∼100 trials on average). If the valve did not deliver the water reward, the session was excluded from the analysis.

#### Sound-only sessions

In these control experiments, the pole was angled forward such that it would not be able to reach the whisker field and be detected by whisking. Mice completed this version of the task for two consecutive days, with regular sessions for monitoring of baseline behavior (Before in Fig.3), followed by two sound-only sessions (Sound-only in Fig 3) and finishing with two regular sessions (After in Fig.3). Sessions from the two days were pooled together by category for analysis.

#### Recall experiments

After the end of the initial training period, mice were progressively allowed to drink water ad libitum and kept from 11 to 18 weeks before re-testing. During recall training, mice were not given help sessions and were tested for 500 trials over multiple days.

### Behavioral data analysis

Behavioral data was analyzed with custom code, written in Matlab. Statistical tests were performed in Matlab using its standard statistical packages (R2021b). Linear-mixed models included individual subjects as random effects, genotype and time (in the appropriate unit, for example session or day) as fixed effects and the variable tested as observations.

#### Licking behavior

The result of each trial was determined based on the trial type presented (Go or NoGo) and the mouse’s actions (Lick or NoLick). Hit and reject rates were calculated as the number of hits (correct Lick) or rejects (correct NoLick) per session divided by the number of Go or NoGo trials respectively. Time spent in timeout was obtained by subtracting total trial duration with first lick time for false alarm trials:

#### Sensitivity

Sensitivity was measured with the discriminability index d’, which was calculated as *d*′ = *Φ*^−1^(ℎ*it rate*) − *Φ*^−1^(*false alarm rate*), where *Φ*^−1^ is the inverse cumulative distribution function of the standard normal distribution.

#### Information

Shannon’s measure of information is related to the amount of surprise or uncertainty associated with a particular event. Mathematically, for a given discrete random variable *X*, it is defined as:

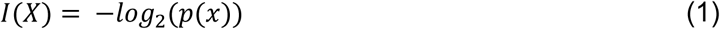

with *p*(*x*) corresponding to the probability of a given event.

The average information, also called entropy (*H*), is the weighted average of the Shannon’s information of all possible events from the variable’s probability distribution

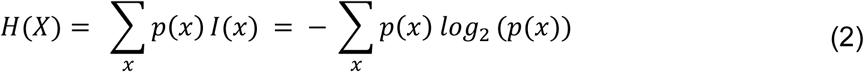

In our task, we consider two variables: stimulus (*S*) and action (*A*). We calculated the entropy *H*(*S*) and *H*(*A*) for both the stimulus and response based on the relative frequencies of trial types (Go/NoGo) and actions (Lick/NoLick) respectively.

Mutual information (*I*) is a measure of dependency of two variables. If mutual information is equal to 0, the two variables are independent, and knowing the value of one variable does not help predict the value of the other. It is calculated from the joint probability distribution of states and actions, p(s,a), as

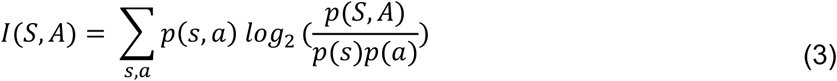

which can be rewritten as,

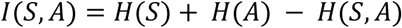

with the joint entropy of S and A:

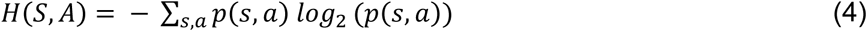

In plots where trials were divided in blocks, the entropies were estimated using the block trial number to have the most accurate estimation of mutual information within that time interval.

Because mutual information can be biased when the number of samples are limited, we randomly shuffled stimulus labels to generate null distributions in the case of no relationship between actions and task conditions. These values were approximately zero, with small positive residuals consistent with finite sample bias (**Supp Fig 1**), although the bias was slightly larger when mutual information was estimated from the first 10 sessions only. Importantly, even during the first 10 sessions, real mutual information values were significantly higher than the shuffled distribution, which had values one order of magnitude smaller on average (**Supp Fig 1**; mean shuffled WT: 0.0031 ± 0.0049 s.d; mean shuffled SRGAP2C: 0.0032 ± 0.0034 s.d, **Fig 4E**; mean WT: 0.019 ± 0.026 s.d, mean SRGAP2C: 0.016 ± 0.019 s.d).

Transmission efficiency is a measure of how efficiently information is communicated across an information channel. It is defined as the proportion of the output entropy that is also shared by the mutual information between input and output, the remainder being due to noise or other unmeasured variables. It was calculated as *Te* = *I*(*S*, *A*) / *H*(*A*) where I (S, A) is the mutual information between states and actions, as defined above, and H(A) is the entropy of the action distribution.

### Reinforcement learning model

The RL model was written with custom code in Julia. Given that licking in this task is the only way to learn stimulus-outcome associations, but with a risk of penalty in terms of average reward obtained (timeouts), we modeled the Go/No-Go task as an infinite-horizon, average-reward Markov Decision Process (MDP) defined by the tuple (S, A, R, T).

#### MDP formulation of the task

**States S.** The model assumes four possible states: inter-trial interval (ITI), stimulus present (SP), stimulus absent (SA), and timeout (TO).

**Actions A.** The possible actions in any of these states are Lick or No Lick.

**Rewards R**. The possible rewards are +1 and 0. The agent receives reward +1 only if the state is SP and the action is Lick.

**Transition T.** The dynamics of the task are very simple: ITI transitions to SP with probability 0.6 and to SA with probability 0.4, regardless of action. SP transitions to ITI regardless of action (Hit if action is Lick, Miss if action is No Lick), and SA transitions to ITI if action is No Lick (Correct rejection), and transitions to TO if action is Lick (False Alarm). Finally, if the agent is in TO and takes action Lick, it goes back to TO.

#### Objective and constraints

To model habit formation and the cost of cognitive control, we introduce an information-theoretic constraint on the agent’s policy. The agent seeks a policy *π*(*s*, *a*) that maximizes the long-run average reward *ρ*, subject to a penalty on the policy complexity, quantified by the mutual information between states and actions *I*(*S*; *A*). The objective is to maximize the unconstrained Lagrangian:

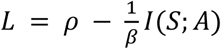

where *β* > 0 is an inverse temperature parameter dictating the trade-off between reward maximization and complexity minimization, we call it the capacity of the agent. As *β* → ∞, the agent acts as a standard reward-maximizer; as *β* → 0, the agent’s policy collapses to its marginal action distribution (its habit), irrespective of the state.

#### Policy and soft value function

To satisfy the complexity constraint, the optimal policy takes the form of a Gibbs distribution biased by a marginal prior *P*(*a*), which represents the agent’s global action frequencies (the habits):

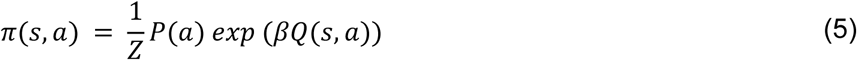

where *Q*(*s*, *a*) is the differential action-value function estimating the expected future reward relative to the average reward *ρ*.

Given that agents pay a complexity price, the resulting differential state-value function V(s) is a soft transformation of *Q*(*s*, *a*), capturing the value of the state under the complexity-constrained policy:

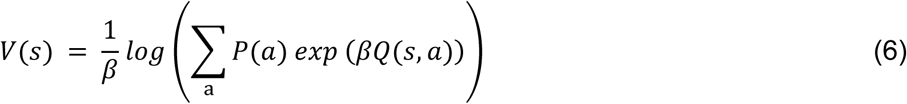

#### Online learning of values and average reward

We approximate the optimal policy using a variant of the differential Q learning algorithm (*78*). At each time step *t*, after taking action *a_t_* in state *s_t_* and receiving reward *r_t_* and transition to *s_t_*_+1_, the agent computes a differential TD error:

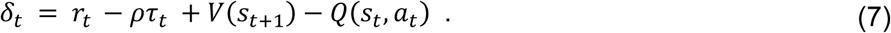

To account for the semi-Markov nature of the task (where different states have varying durations, such as ITI or timeouts), the average reward penalty is scaled by the time elapsed *τ_t_*.

The state-action values and the global average reward estimate are updated via stochastic gradient descent:

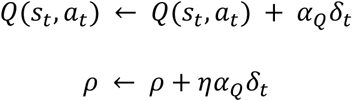

where *α_Q_* is the learning rate for the Q-values and *η* is the scale of learning rate for average reward compared to the one for Q-values.

#### Online habit formation (marginal prior update)

The optimal marginal prior *P*(*a*) is defined as the stationary distribution of actions marginalized over the state visitation frequencies. To compute this online without explicit knowledge of the state transition dynamics, we utilize an Exponential Moving Average (EMA) that incrementally shifts the prior toward the most recently chosen action:

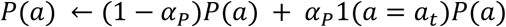

where 1() is the indicator function and *α_P_* is a slow learning rate *α_P_* < *α_Q_*. This separation of timescales ensures that the habit tracks the slowly evolving stationary policy without destabilizing the fast convergence of the differential Q-values.

#### Parameters

The model has some free parameters, 3 related to the speed of learning, and 1 for the capacity of the agent. To obtain the results from Fig. 5, we fix all the learning rates to obtain qualitatively similar results to experimental data in terms of average reward. Specifically, *α_Q_* = 0.02, *α_P_* = 0.01, *η* = 5. Two populations 40 agents differ only in their capacity, *β_low_* = 1.0, *β*_ℎ*ig*ℎ_ = 1.7, which were chosen to reproduce qualitatively the mutual information obtained by the policies of the two groups. The Q-values were initialized to zero, and the habits were initialized around 0.5 for each action. We inject some variability to avoid artifacts due to an arbitrary initial choice of habits, so each agent was initialized with a uniformly random P(lick) between 0.4 and 0.6.

#### Relationship to prior work

Our algorithmic approach builds upon the policy compression framework introduced by Lai and Gershman (*47*). While their framework shares the fundamental mathematical objective of maximizing reward under a mutual information *I*(*S*; *A*) constraint, our specific learning rules differ from theirs in two critical ways to accommodate our experimental paradigm:

1. Value-based vs. actor-critic: Lai and Gershman propose an actor-critic (policy gradient) architecture. In their formulation, the complexity cost (the log-ratio log 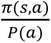) is explicitly subtracted from the reward signal to formulate the prediction error, and a separate parameterized actor is updated via gradient ascent. In contrast, our approach is strictly value-based. We handle the complexity constraint implicitly within the softmax policy and the state-value calculation, completely eliminating the need for a separate parameterized actor and providing faster, stable convergence in tabular state spaces.
2. Sequential MDPs vs. contextual bandits: The online learning rules derived in the appendix of Lai and Gershman (2021) are designed for tasks without sequential state dependencies (effectively contextual bandits), utilizing a target based on the immediate Reward Prediction Error (RPE). Because the Go/No-Go task includes time-delayed consequences (e.g., timeouts), we generalized their online policy compression rules to the infinite-horizon MDP setting by replacing the myopic RPE with a differential temporal difference error (TDE), ensuring the algorithm accounts for both future states and the average rate of reward.

## ACKNOWLEDGEMENTS

This work is dedicated to the memory of co-author Roberto Araya (1974–2024). The authors would like to thank members of the Araya, Di Cristo and Ebitz labs, Bénédicte Amilhon and Christopher Pack for useful discussions and advice. This project was funded by the Canadian Institute of Health Research (MOP-202203; R.A), a Natural Sciences and Engineering Research Council of Canada (NSERC Discovery Grant, RGPIN-2022-04275; R.A), the Air Force Office of Scientific Research (FA9550-24-1-0305; R.B.E.), NSERC Discovery Grant (RGPIN-2026-05793; R.B.E.), the Canada Research Chair in the Dynamics of Cognition (FD507106; R.B.E.), two recruitment scholarships from the Neuroscience department, Université de Montréal (C.S.B), the Roberto Araya scholarship for Excellence in Neuroscience Research (C.S.B), and the Energir Scholarship for Excellence in Research (C.S.B) from the CHUSJ foundation, the National Institute of Health-National Institute of Neurological Disorders and Stroke-R35 NS127232 (F.P) and an award from the Nomis Foundation (F.P). This material is based upon work supported by the Air Force Office of Scientific Research under award number FA9550-24-1-0305. Any opinions, findings and conclusions or recommendations expressed in this material are those of the author(s) and do not necessarily reflect the views of the United States Air Force.

## Supplementary figures

**Supplementary figure 1.**
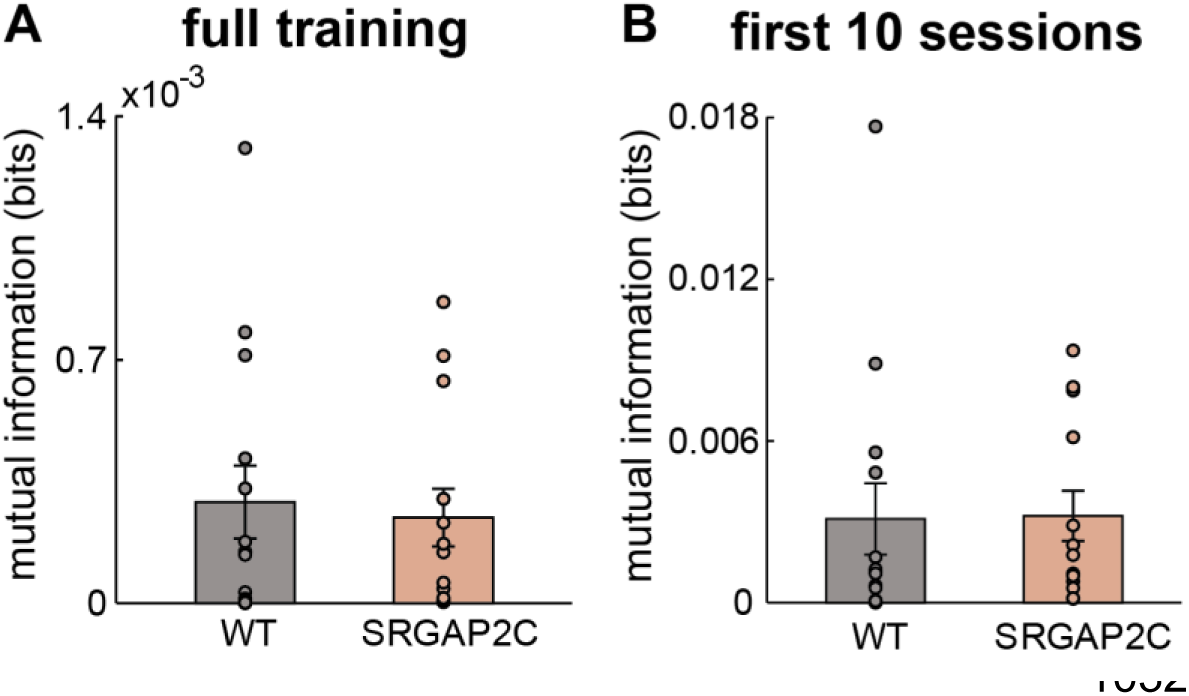
Mutual information with shuffled stimulus labels. WT (n=14) and SRGAP2C (n=13) are shown in grey and pink beige respectively. A. Shuffled mutual information values for the full training epoch. Significance was tested with a Wilcoxon ranksum test (mean WT: 0.00029 ± 0.0004 s.d; mean SRGAP2C: 0.00025 ± 0.0003 s.d, *p* = 0.9). B. Shuffled mutual information values for the first 10 sessions. Significance was tested with a Wilcoxon ranksum test (mean WT: 0.0031 ± 0.0049 s.d; mean SRGAP2C: 0.0032 ± 0.0034 s.d, *p* = 0.4)

**Supplementary figure 2.**
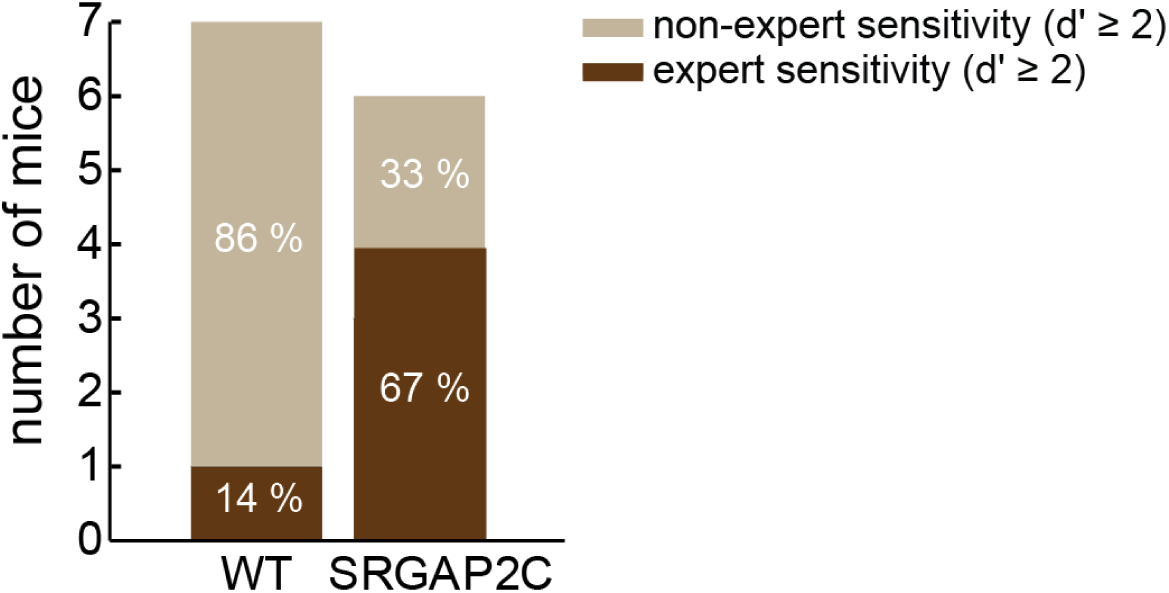
Sensitivity during recall. A subset of mice (WT: n=7; SRGAP2C: n=6) were re-exposed to the task after 11-18 weeks of ad libitum access to water and normal cage activity. Number of mice showing expert sensitivity (d’ ≥ 2) across genotypes. A higher group of SRGAP2C mice (4/6, 67%) reached the threshold sensitivity compared to WT mice (1/7, 14%); however this difference did not reach statistical significance (Fisher’s exact test: *p* = 0.1).

